# Treerecs: an integrated phylogenetic tool, from sequences to reconciliations

**DOI:** 10.1101/782946

**Authors:** Nicolas Comte, Benoit Morel, Damir Hasic, Laurent Guéguen, Bastien Boussau, Vincent Daubin, Simon Penel, Celine Scornavacca, Manolo Gouy, Alexandros Stamatakis, Eric Tannier, David P. Parsons

## Abstract

**Motivation:** Gene and species tree reconciliation methods are used to interpret gene trees, root them and correct uncertainties that are due to scarcity of signal in multiple sequence alignments. So far, reconciliation tools have not been integrated in standard phylogenetic software and they either lack performance on certain functions, or usability for biologists.

**Results:** We present Treerecs, a phylogenetic software based on duplication-loss reconciliation. Treerecs is simple to install and to use. It is fast and versatile, has a graphic output, and can be used along with methods for phylogenetic inference on multiple alignments like PLL and Seaview.

**Availability:** Treerecs is open-source. Its source code (C++, AGPLv3) and manuals are available from https://project.inria.fr/treerecs/

## Context

Phylogenetic reconciliation methods are recognized to be powerful tools to understand the evolution of gene families (Szöll ő si *et al*., 2015), and host-parasite co-evolution (Bailly-Bechet and *et al*, 2017). They consist in annotating, rooting or improving gene (parasite) trees by comparing them to species (host) trees, under evolution models accounting for events such as duplications, transfers, losses, and incomplete lineage sorting. Available tools (e.g. Bansal and *et al*, 2018; Stolzer and *et al*, 2012; Jacox and *et al*, 2016; Akerborg and *et al*, 2009; Szöllő si and *et al*, 2013; Morel *et al*., 2020) take into account duplications, transfers, losses or incomplete lineage sorting, with binary or multifurcated gene or species trees. They either output one solution, optimizing a parsimony score or a likelihood function, or sample according to a likelihood distribution. Yet, several reconciliation tools rely on external libraries making the installation tedious and difficult. Moreover, the formats for gene and species names, reconciliations and reconciled gene trees are often very specific, not flexible, and not compatible with one another. Finally, basic functionalities are missing, e.g. no current software can efficiently correct and root multifurcated gene trees at the same time.

The aim of the new software we present here, Treerecs, is to bring reconciliation tools to the level of usability of standard phylogenetic tools and interface with them for complete biological analyses, from sequence alignment to reconciliation, via a graphical interface and graphical output.

## Usability of Treerecs

Treerecs is available on Linux, Mac OSX and Microsoft Windows, and does not require any external library to be used^1^. Debian and RPM packages are currently in preparation.

As do all reconciliation software, Treerecs requires 3 kinds of information: a rooted binary species tree, one or more gene trees (rooted or not, binary or not) and a mapping between the leaves of gene trees and those of the species tree. One frequent difficulty with reconciliation software is the strict formats for the inputs. Treerecs accepts Newick, NHX and PhyloXML, and does not require any special treatment for special characters frequently reserved for the format (*e*.*g*. @#_|) that may also be present inside gene or species names. The mapping can be informed by the user in a separate file, with a versatile format. Alternatively, the mapped species can be specified in the gene names. In that case, the species name corresponding to a given gene is directly sought in its name, with no requirement on its position nor on the separation character. Unless there is an ambiguity, *i*.*e*. if the gene name contains several species names, Treerecs is always able to infer the mapping automatically.

The output can also be given in a variety of standard formats, including RecPhyloXML, shared by several other tools for reconciled gene trees (Duchemin and *et al*, 2018), and SVG for display purposes (see Figure 1a).

**Fig. 1.**
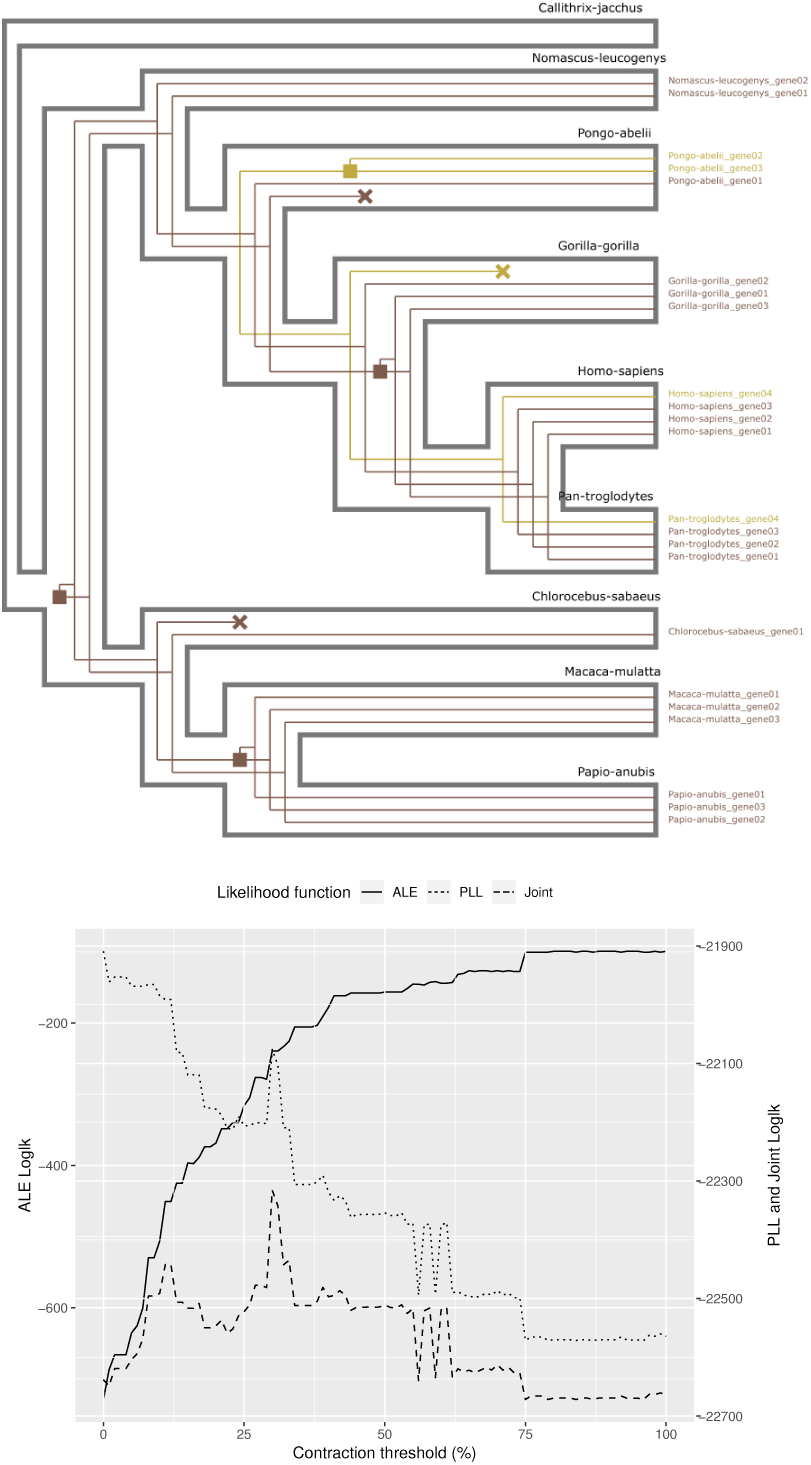
(A) Example of SVG output produced by Treerecs. Treerecs can draw several gene trees (two here) inside the associated species tree. Squares represent gene duplications events and crosses, losses. (B) Likelihoods of ALE (on reconciliation) and PLL (on sequence information) have respectively an increasing and decreasing trend, function of the contraction threshold. The joint likelihood has its maximum at 31%.

## Reconciliation, Rooting, Resolution, Correction

Basic functionalities of Treerecs include: (i) computing and depicting reconciled gene phylogenies within the associated species tree with duplications and losses, (ii) rooting gene trees by identifying a root that minimizes the duplication and loss score (iii) correcting gene trees by contracting weakly supported branches (according to a contraction threshold) (iv) resolving multifurcations (either given as an input or being the result of contractions) minimizing the duplication and loss score. Resolution and correction are achieved using the ProfileNJ algorithm (Noutahi and *et al*, 2016). Note that the resolution and rooting can be done at the same time, a feature no other current reconciliation software achieves, except ecceTERA which has an exponential running time on correction part (but also accounts for gene transfers). When there are several solutions for rooting, resolving or correcting, Treerecs outputs randomly one or more solutions from a uniform distribution. We provide metrics (likelihoods computed from site substitution models and from gene content models) to give the possibility to choose among these solutions.

## Integration

Treerecs offers more than just the basic functions presented above. In particular, it can compute the phylogenetic likelihood of a tree given a multiple sequence alignment, using the Phylogenetic Likelihood Library (PLL — Flouri and *et al* (2014)). It can also compute the reconciliation likelihood of a gene tree given the species tree with the ALE DTL model (Szöll ő si and *et al*, 2013). This allows, for instance, to explore the gene tree space, via a variation of the contraction threshold for branches with low support, scored by a joint likelihood. This procedure is illustrated in Figure 1b and in the next section.

Furthermore, Treerecs is integrated in the current version 5.0 of Seaview (Gouy *et al*., 2020), already available from the Treerecs website.

Thus, it can be used with a graphical interface along with a standard phylogenetic pipeline.

## Efficiency

Treerecs accomplishes comparable tasks with Ranger-DTL (Bansal and *et al*, 2018), Notung (Stolzer and *et al*, 2012) and ecceTERA (Jacox and *et al*, 2016). We tested the efficiency of all of them on their basic functions (rooting, correction, rooting+ correction), on trees with variable sizes from the Ensembl Compara database (Herrero *et al*., 2016). With the exception of the rooting task, for which ecceTERA is notably the best, Treerecs is computationally more efficient than competing tools. We report the performance of the “correction + rooting” task in Table 1, along with a comparison of different features.

**Table 1.**
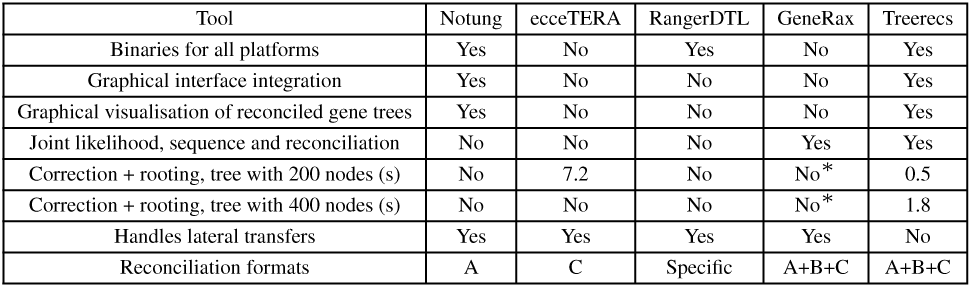
Novelties and efficiency of Treerecs, by comparison with Notung, ecceTERA, GeneRax and RangerDTL, on several features. In the last row, A, B and C stay respectively for NHX, Newick and RecphyloXML. (*) GeneRax achieves a sort of gene tree correction and rooting in decent time but the comparison is difficult because the problems solved by Treerecs and GeneRax are different (GeneRax optimises according to the joint likelihood function). Another tool we would like to mention here is TreeFix-DTL (Bansal et al., 2015), which would exhibit the same column entries than RangerDTL in the table, but has the additional feature to combine the information of the sequence alignment with the cost of the reconciliation: only gene trees that are statistically equivalent to the input gene tree are considered when correcting it by minimizing the duplication and loss score.

To assess the efficiency of the joint likelihood principle to recover gene trees, we simulated 656 gene trees with Zombi (Davín and *et al*, 2019). Zombi can jointly simulate a species tree (with the possibility that some species go extinct, 26 remained in our analysis), and genomes (genes, gene order, sequences). The average normalized Robinson-Foulds (RF) distance of inferred trees to true trees is 0.167 for trees inferred by RaxML and 0.117 for trees infered by Treerecs (optimizing the joint likelihood). We found that selecting the contraction threshold that maximizes the joint likelihood gives a better average RF distance to the true trees than arbitrary choosing any of the possible thresholds. This shows, first, that reconciliations can be a good guide and help finding better trees than methods based on sequences only, and second, that the best approach combines the two likelihoods (data available on the Treerecs website).

## Footnotes

1 CMake and a C++14-apt compiler are required to *compile* Treerecs.

